# Linking land□use change, water quality, and host-parasite dynamics with droplet digital PCR and Bayesian path analyses

**DOI:** 10.64898/2026.05.12.724588

**Authors:** Ipsita Srinivas, Chloe A. Fouilloux, John Berini, Paul Orlando-Simoni, Eric Neeno-Eckwall, Heather Alexander, Emma Choi, Grace Vaziri, Amy Chen, Gwen Casey, Shira Dubin, Cate Patterson, Amanda K. Hund, Daniel I. Bolnick, Jessica L. Hite

## Abstract

Global changes in land use and nutrient cycling are transforming ecosystems at unprecedented rates, with significant consequences for infectious disease dynamics. Aquatic environments are particularly vulnerable because the interplay of habitat modification, nutrient enrichment, and biodiversity loss can drive pronounced changes in the community composition of food webs, including hosts and parasites. Yet, despite well-documented effects of habitat modification on aquatic communities and food webs, the mechanisms through which these changes influence infectious disease dynamics remain poorly resolved. This gap arises, in part, because it remains challenging to disentangle how multiple stressors interact to shape disease outcomes and quantify parasite levels and host densities from field-collected samples. Here, we illustrate two tools that might help address these challenges. First, highly sensitive droplet digital PCR can quantify infection loads even when the signal:noise ratio is low. Second, stepwise Bayesian path analyses can identify the direct and indirect pathways connecting land-use changes to infectious disease dynamics. As a case study, we examined cyclopoid copepods and their helminth parasite, *Schistocephalus solidus*, across 47 freshwater lakes on Vancouver Island, a region strongly shaped by commercial logging, including widespread clear-cutting of old-growth forests. Our results reveal a positive correlation between copepod density and deforestation, potentially mediated by associated changes in water quality and calanoid copepods, key competitors of the focal host. ddPCR enabled sensitive detection of extremely low parasite signals in field-collected copepods. We detected positive infections in only 19.5% of the lakes surveyed, highlighting the difficulty of assessing disease dynamics in natural populations. Nonetheless, this study highlights the challenges of linking land-use change to disease outcomes, while also demonstrating that sensitive molecular and statistical tools offer new ways to reveal these hidden connections.

## Introduction

Human-driven changes to landscapes are reshaping ecosystems worldwide, with profound consequences for the emergence and transmission of infectious diseases^1,2^. Aquatic ecosystems are especially sensitive to these stressors, as land-use change, nutrient enrichment, and biodiversity loss interact to alter community structure and, subsequently, disease dynamics^3–13^. Deforestation, in particular, can alter water quality in aquatic systems through increased allochthonous nutrient inputs, altering resource availability, with cascading effects on food-web dynamics, including shifts in the abundance, distribution, and diversity of hosts and parasites^4,9,14–20^. Understanding the mechanisms by which shifts in habitat modulate the population dynamics of hosts and parasites is crucial for identifying ecological levers that can be managed to reduce infection risk, anticipate emerging threats, and guide conservation and public health interventions^21^. Yet, despite clear habitat-mediated effects on aquatic ecosystems and food webs, causally linking these changes to host-parasite dynamics or infectious disease outcomes remains challenging.

This gap in knowledge is particularly acute for the numerous macroparasites with complex life cycles such as helminths, including roundworms, tapeworms, and flukes. These parasites remain a persistent challenge across public health, wildlife, and livestock systems, and many require aquatic invertebrates as intermediate hosts^22,23^. Given their critical role in the transmission of parasites with complex lifecycles, intermediate hosts can act as transmission bottlenecks but are notoriously challenging to study^24^. Copepods - aquatic crustaceans - exemplify this challenge. Because copepods are small-bodied, cold-blooded, omnivorous consumers and predators near the base of aquatic food webs, they are particularly sensitive to environmental changes, such as fluctuations in temperature and primary production^11,25–30^.

These ubiquitous zooplankton serve as the first intermediate host for many macro and micro-parasites affecting wildlife, livestock, and humans^31^. Notable examples of helminths transmitted by copepods include *Gnathostoma* and *Spirometra* spp., whose larval stages are responsible for human larval migrans, or ’creeping eruptions’^22^. Similarly, helminths in the genus *Dracunculus*, which cause debilitating pain and disability in humans and wildlife. Despite extensive control efforts dramatically reducing infection numbers from 3.5 million in 1986 to 27 human cases in 2020, eradication remains challenging due to recent discovery of, and increases in, non-human hosts - in part due to the difficulties in monitoring and controlling the intermediate copepod host^32,33^.

Moreover, mounting evidence highlights that for many parasites with complex life cycles, like helminths, environmental drivers often act indirectly by restructuring the density and distribution of intermediate host communities rather than acting directly on the parasites themselves^5,34,35^. Changes in host density can cause cascading, non-linear effects on parasite transmission in definitive hosts, many of which modulate spillover to livestock and humans^35,36^. Consequently, strong environmental gradients do not always produce predictable patterns of infection in many host-parasite systems, but especially in helminths with complex life cycles. From an applied perspective, overlooking the intermediate hosts could dimmish the efficacy of other control efforts and even backfire, leading to larger and more severe outbreaks^37^.

Macroparasite systems involving copepod hosts may be especially sensitive to environmental disturbances because even modest shifts in copepod density can drive pronounced changes in transmission. For example, seasonal transmission of the helminth *Dracunculus medinensis* increases during hot, dry periods when receding water levels create shallow ponds that support higher densities of copepod hosts, thereby elevating exposure risk for humans^38,39^. Beyond temperature, copepods are highly sensitive to shifts in water quality driven by nutrient enrichment, acidification, and changes in water temperature^29,40–43^. Indeed, given their importance in both marine and freshwater aquatic food webs, and their pronounced responses to environmental variation, copepods are frequently used as indicators of ecological change^41,43,44^. This raises an important question: could copepods also serve as indicators of disease risk for macroparasites in increasingly human⍰modified ecosystems?

Answering this question carries important implications for both conservation and public health. Helminth⍰borne diseases remain historically understudied, despite infecting 1.5 billion people worldwide, particularly in regions of the world that are also experiencing rapid climate and land⍰use change^45^. Identifying and interrupting the ecological drivers of transmission is therefore a critical research priority. However, despite the central role that copepods play in numerous helminth life cycles, detecting or quantifying infection in these small hosts remains notoriously difficult^31^. As a result, most studies focus on vertebrate hosts, leaving the ecological factors regulating parasite dynamics within copepod populations largely overlooked.

Community ecology provides a powerful framework for linking such habitat changes to the population dynamics of hosts and parasites by tracing how environmental variation restructures host communities, their prey, key competitors, and associated parasites. In multi⍰host, multi⍰trophic systems, however, these pathways are often difficult to resolve because environmental change can strongly influence host density and community structure without necessarily producing linear or predictable responses in infection intensity or prevalence^36^. This challenge is especially pronounced when parasites occur at low densities or are patchily distributed, making it difficult to detect consistent infection signals.

Here, we highlight two complementary approaches to address these challenges. First, we use highly sensitive droplet digital PCR to quantify helminth infection loads within copepod hosts, even when the signal:noise ratio is low. Second, we evaluate theoretically motivated and empirically supported relationships between environmental variables and copepod dynamics (Table 1) using univariate field patterns (see SI Text 3; SI Figures S1-S4). Then, we apply step-wise Bayesian analyses to identify the direct and indirect pathways connecting land-use changes to shifts in host density.

**Table 1:**
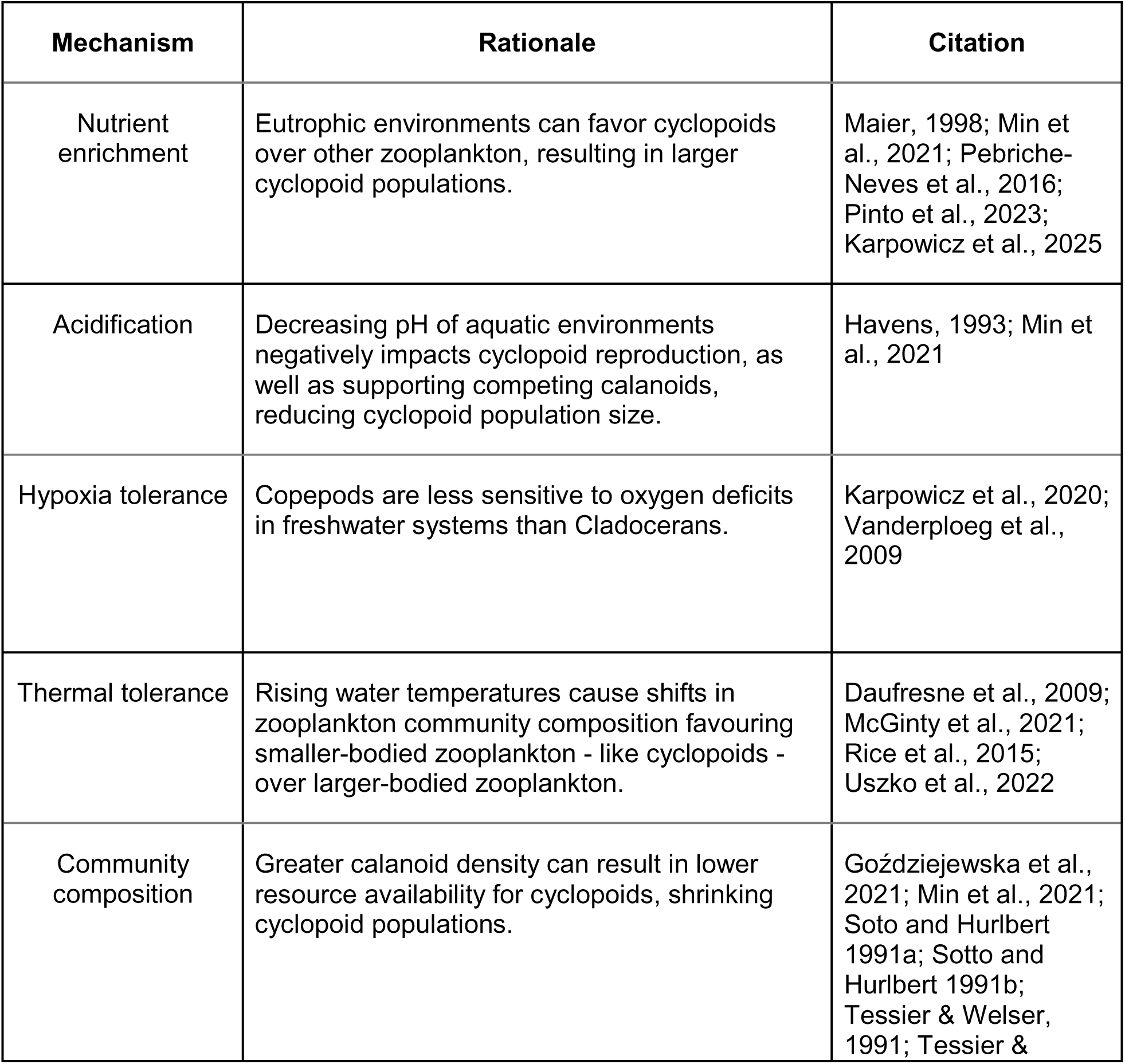
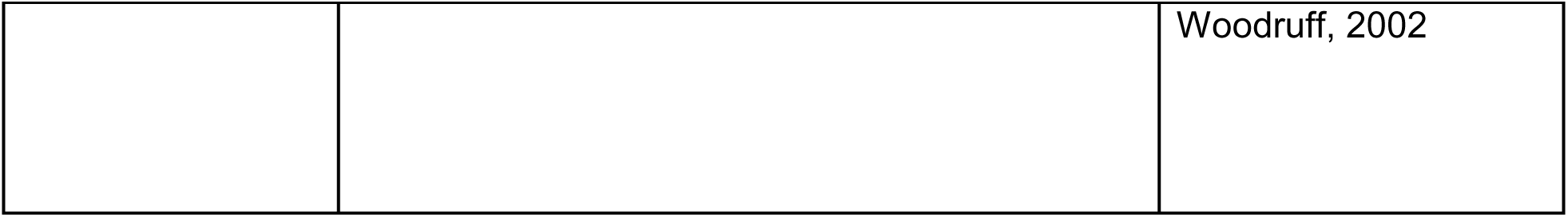
Hypothetical key drivers of spatial variation in cyclopoid copepod abundance.

As a case study, we focus on a well⍰characterized freshwater parasite system involving Schistocephalus solidus, a helminth that uses copepods as its first intermediate host. This trophically transmitted parasite has a complex life cycle that spans three hosts across different trophic levels: cyclopoid copepods, the threespine stickleback (Gasterosteus aculeatus), and fish-eating birds (Fig. 1)^46^. In this system, copepods are a critical component in the life cycle of S. solidus, making them a sensitive point of control for trophic transmission to successive hosts. Previous studies, however, have largely focused on interactions between the focal parasite and threespine stickleback^47–55^, leaving key questions^31^ open concerning the copepod hosts^46,56–64^.

**Figure 1:**
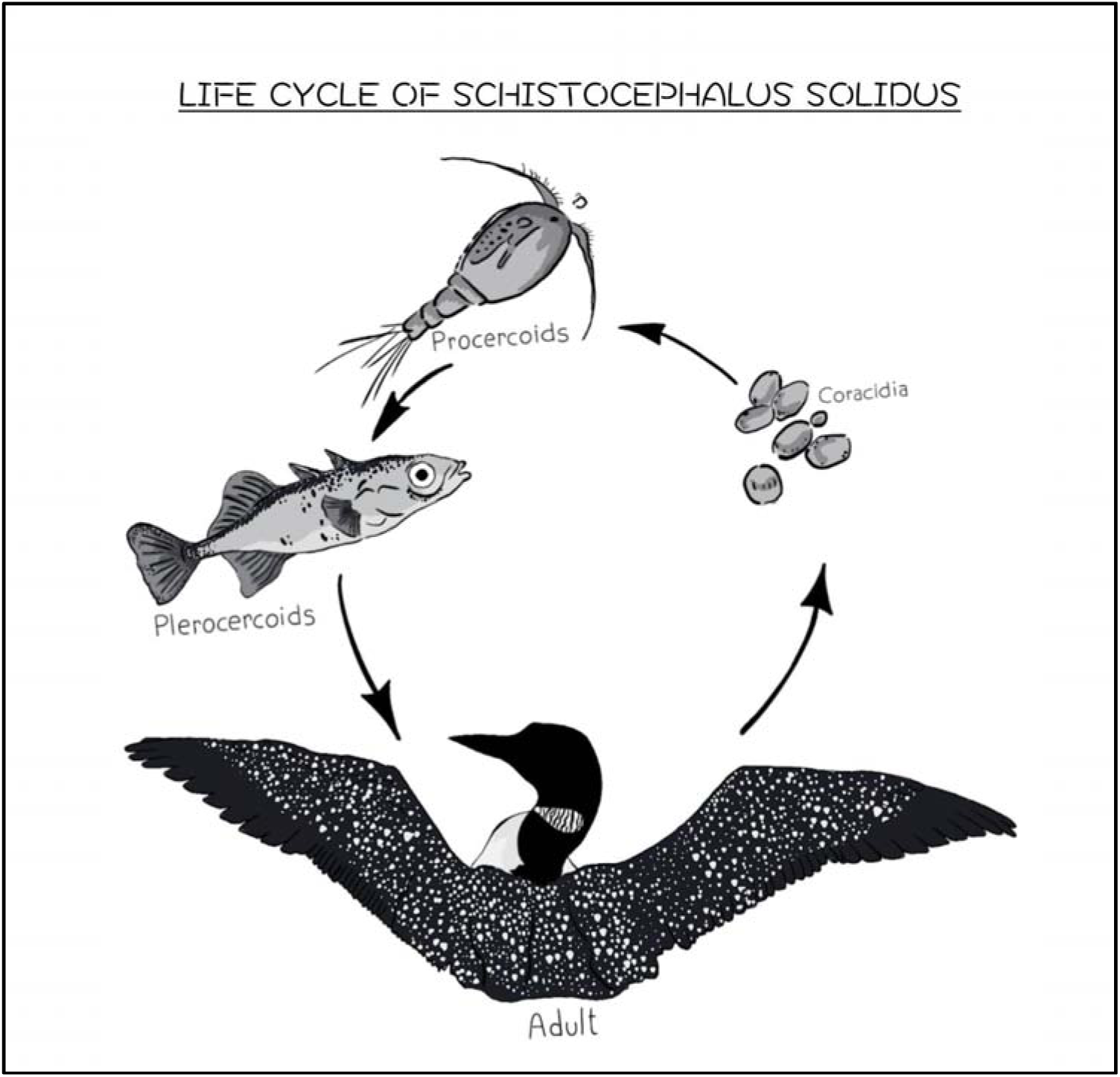
Life cycle of the helminth *Schistocephalus solidus* involving cyclopoid copepods, stickleback fish, and piscivorous birds. Cyclopoid copepods serve as the first intermediate host, becoming infected when they ingest free□swimming coracidia hatched from eggs shed in bird feces. Within 9-14 days, parasites develop into procercoids inside the copepod. When an infected copepod is consumed by a threelJspined stickleback, the second intermediate host, the parasite penetrates the gut wall and grows into a plerocercoid in the body cavity. These plerocercoids can reach large sizes and manipulate host behavior, increasing the fish’s susceptibility to predation by loons or herons. Once consumed by a piscivorous bird, the parasite rapidly matures, reproduces, and releases eggs back into freshwater, completing the cycle. Infection load in copepods depends on encounter rates with coracidia and copepod density. Figure credit: Raquel Santamaria Germani; Amanda K. Hund.

Our primary goal is to identify the environmental pathways that regulate host availability, because host density and community composition fundamentally constrain opportunities for parasite transmission. To do this, we quantify how landscape disturbance and land-use mediated changes to water quality shape both cyclopoid hosts and calanoid copepods, which are both prey and key competitors with cyclopoids, but are not known to become infected with S. solidus. This focus allows us to evaluate the mechanisms by which habitat alteration propagates through the zooplankton community, mechanisms that ultimately set the upper limits on parasite encounter and infection, even when infection patterns themselves are difficult to detect or highly variable.

Our work centers on freshwater lakes across Vancouver Island, a region that supports extensive commercial logging, including widespread clearcutting of old⍰growth forests. These anthropogenic activities can substantially reshape aquatic ecosystems by altering nutrient inputs, sedimentation, and shifts in water quality, including eutrophication in more extreme cases^3,17,18,65^. We hypothesized that percent forest loss would alter water quality via increases in allochthonous nutrient and organic matter inputs, which can stimulate primary production and lower pH through the delivery of humic and organic acids. Increased ecosystem respiration from both allochthonous and autochthonous organic matter may reduce dissolved oxygen, especially deeper in the water column. Further, reduced riparian shading should increase water temperature. Each of these factors can strongly influence the density and community composition of copepods.

Across lakes, we find evidence that deforestation is linked to shifts in copepod density and community structure, and that these biotic responses correspond with land⍰use–associated gradients in nutrients and water⍰quality variables. ddPCR enabled sensitive detection of extremely low parasite signals in field-collected copepods. Nonetheless, we detected positive infections in only 19.5% of the lakes surveyed. As a result, relationships between infection load and environmental metrics remain weak, even in systems where copepod responses to environmental change are strong. Together, our findings illustrate both the challenges (and opportunities) of tracing mechanistic pathways from habitat alteration to disease.

## Methods

### Focal Host and Parasite

Our focal host, cyclopoid copepods, are selective grazers in freshwater lakes across North America, including the lakes on Vancouver Island studied here^66–68^. In many lakes, cyclopoid copepods specifically are hosts to a parasitic tapeworm, Schistocephalus solidus (hereafter referred to as “tapeworm”). This trophically transmitted parasite has a complex life cycle that spans three hosts across different trophic levels: cyclopoid copepods, the threespine stickleback (Gasterosteus aculeatus), and fish-eating birds^48^. Successful development and reproduction require the parasite to infect each of these hosts sequentially. The cycle begins in freshwater, where eggs released from the definitive avian host hatch into free-swimming larvae known as coracidia. These larvae are consumed by cyclopoid copepods, the first intermediate host. Within 9-14 days, the coracidia develop into procercoid larvae inside the copepod^46,69^.

When an infected copepod is eaten by a threespine stickleback, the second intermediate host, the procercoid penetrates the intestinal wall and migrates into the fish’s body cavity. There, it develops into a plerocercoid over several weeks to months. During this stage, the parasite can grow to occupy a substantial portion of the host’s body cavity, often reaching up to half the host’s body mass. Infected sticklebacks exhibit altered behaviors that increase their susceptibility to predation by piscivorous birds, such as loons or herons. Once consumed by a bird, the plerocercoid rapidly matures into an adult tapeworm in the bird’s gut. Mating occurs shortly thereafter, and fertilized eggs are released into the environment via the bird’s feces, completing the cycle^48,70^. With this natural history, transmission and therefore infection load in copepod hosts could increase through density-dependent transmission and thus, higher copepod densities or higher density of coracidia^71^.

### Field Surveys

To examine the abiotic and biotic factors driving spatial variation in host-parasite dynamics of the focal copepod-helminth system, we sampled 47 lakes across Vancouver Island, B.C., Canada (Fig. 2), during the breeding season of the intermediate copepod host (June 2023). The first intermediate copepod host is present in all sampled lakes. At each lake, we collected zooplankton community composition data, limnological, and bathymetric data using previously established methods^72,73^. We compiled variables related to land cover, forest structure, and land use, as well as proximity to human activity and infrastructure, from the iMapBC interface, except for NDVI and EVI indices, which we obtained from NASA’s EarthData platform^74^. We summarized all spatial data within a 1,000 m buffer surrounding each lake, calculating mean, minimum, and maximum values for each covariate. Percent forest loss (% canopy openness) was calculated as 100 − mean canopy (crown) cover (%), constrained to [0, 100].

**Figure 2:**
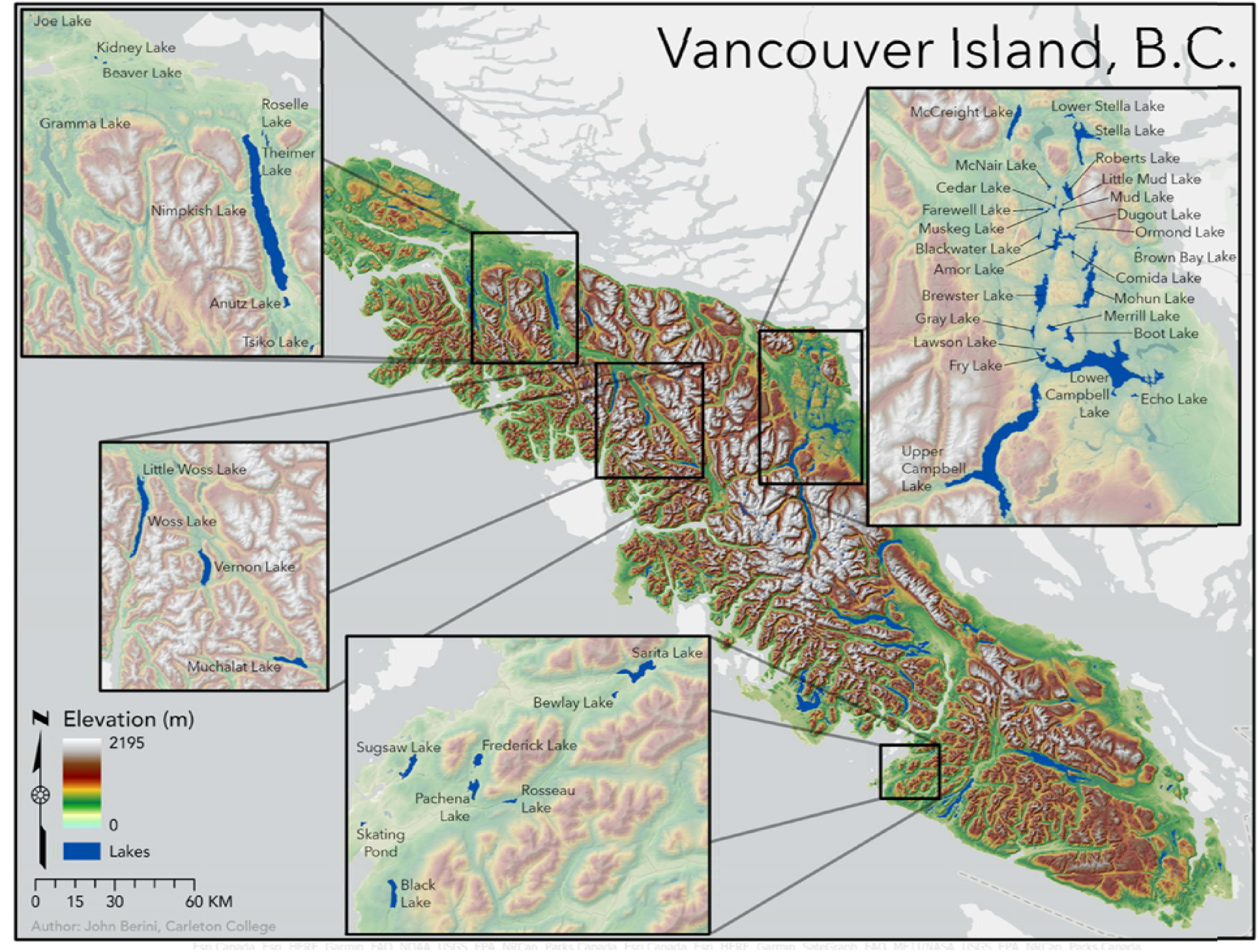
Map of 47-sampled lakes on Vancouver Island, British Columbia. Dark blue shading represents lake locations. Map elevation ranges from 0 - 2195 (m), and is represented by a colour gradient from light green (low elevation) to white (high elevation). Due to sampling and logistical challenges, only 41 of the 47 lakes contained both limnological and zooplankton data and were retained for subsequent analyses. From the 41 lakes retained post-consolidation across datasets, we excluded two lakes identified *a priori* as non-comparable due to depth (Nimpkish > 320m; Muskeg < 2m), leaving 39 lakes. Of the 39 lakes, 28 had deforestation data available after accounting for exclusions and inaccessible proprietary data. Map credit: John Berini.

To examine copepod abundance and zooplankton community composition for each lake, we collected two samples of zooplankton from different locations in the lake, each with three bottom to surface tows of a Wisconsin Net (13 cm diameter, 63-μm mesh). We preserved both samples in 80% ETOH and transported the ethanol-stored samples to the University of Wisconsin, Madison, where they were stored at -4°C. We used the first sample to estimate aerial densities of the focal host (cyclopoid copepods) and all zooplankton taxa within the community (see SI Text 1). Finally, we calculated the inverse Simpson’s diversity index for the total zooplankton community, including both cyclopoid copepods (adult and juveniles, excluding nauplii) and all other co-occurring taxa.

To quantify infection burden in the cyclopoid hosts from the second zooplankton sample, we used droplet digital PCR (ddPCR) following protocols published in Fouilloux et al. (2025). This high-throughput method enables rapid and sensitive quantification of infection load at the population level. Theory predicts that infection dynamics are regulated by population-level processes in host-parasite systems (e.g., host and parasite density)^75,76^. Thus, population-level infection metrics are key to assessing long-term host-parasite dynamics^71,75^. While individual-level assessment of copepod infections would provide direct measures of infection prevalence (the fraction of hosts infected out of the total hosts sampled), the particulars of this system make this logistically challenging. More specifically, visually identifying infected copepods from live or preserved samples was both time intensive and destructive, which would prevent the preservation of samples for future analyses. Given these constraints, we focus here on ddPCR analyses to quantify within-host infection burdens.

Prior to ddPCR, DNA was extracted from zooplankton samples using PowerFecal Pro Kits (Qiagen) with an added 65°C lysis step, quantified spectrophotometrically, and stored at −20 °C. DNA used for ddPCR came from 10 ng µL⁻¹ aliquots of extracts and was pre-diluted (1 ng µL⁻¹; 0.1 ng µL⁻¹) to ensure adequate droplet separation for quantification. Parasite and host DNA were quantified using species-specific 18S rRNA probe-primer sets. Both multiplex and singleplex reactions were run per sample to detect host and parasite DNA while ensuring that multiplexing did not bias quantification or produce false positives. Parasite probes were labeled with FAM and host probes with HEX. A modified universal primer (F517, shortened by two nucleotides at the 5′ end) was paired with parasite-specific platyhelminth primers, and host detection used a cyclopoid-specific probe-primer.

ddPCR was performed on a Bio-Rad QX200 system. Absolute target gene copy numbers were estimated using QuantaSoft software based on Poisson statistics. Negative controls (no template) were included to check for contamination. Population-level infection metrics were calculated by comparing host and parasite copy numbers in each sample to a 1:1 infection standard of a 100 adult copepods, each infected with a single S. solidus. The 1:1 standard also served as our positive control. Wells with fewer than 10,000 accepted droplets were excluded.

To collect the limnological data for each lake, we used an EXO2 multi-probe sonde (YSI Incorporated, Ohio, USA) to measure temperature-depth profiles at one-meter increments to one meter above the hypolimnion. We also collected multiple water quality metrics, including pH, dissolved oxygen (mg L^-1^; also calculated as % saturation), and multiple metrics indicative of nutrient enrichment: phycocyanin, chlorophyll a (a proxy for phytoplankton biomass), and fluorescent dissolved organic matter (fDOM). Measurements for phytoplankton provide insight into qualitative differences among lakes and are units of relative fluorescent units (RFUs). With these data, we calculated the depth of the thermocline, which was then used to calculate refuge size. Refuge size was calculated as the difference between the depth of the thermocline (upper bound, defined as maximum buoyancy frequency) and the oxygen threshold (lower bound, 1mg L^-1^)^77^.

We obtained lake bathymetric data using iMapBC, an interactive mapping platform maintained by the Government of British Columbia. To extract bathymetry data for each of the focal lakes, we applied specific map layers within the platform, including Bathymetric – 7.5M (under “Base Maps”), and Lake Bathymetric Maps and Digital Bathymetric Maps (under “Fish, Wildlife, and Plant Species”). These layers provided standardized measurements for each sampled lake, including perimeter (m), maximum depth (m), mean depth (m), surface area (ha), and elevation (m). We used this data to calculate the Depth ratio, an index of lake basin shape, for each site.

### Statistical Analyses

We examined relationships between deforestation, water quality, copepod density, and parasite infection using both simple univariate analyses and piecewise Bayesian structural equation models (SEM). This SEM approach is particularly well-suited for integration of multiple data streams that inherently vary in completeness.

For example, in our study, bathymetry data were available for all 47 lakes, providing a strong foundation for spatial analysis. Deforestation data, however, were accessible for only 30 lakes, as data sharing by logging companies is voluntary. Additionally, while limnological and zooplankton samples were collected from all 47 lakes, we a priori excluded two lakes because they were too deep (> 300m, Nimpkish) or too shallow (< 2m, Muskeg) to collect representative limnological data with our sonde equipment. Additionally, sonde data from one lake (Dugout) and zooplankton samples from six lakes (Dugout, Bewlay, Frederick, Littlewoss, Sarita, Skating Pond) were lost or damaged during shipping. Thus, some lakes that contained limnological data did not have the accompanying zooplankton data. As a result, post data consolidation and a priori exclusions, 39 lakes included both limnological and zooplankton data, while a subset of 28 lakes had complete data across deforestation, water quality, and zooplankton sampling.

The piece-wise compilation of models in SEM allows us to maximize the use of available data, maintaining analytical power for pathways that do not depend on specific datasets, while still enabling robust evaluation of land-use effects where data are present.

First, as an initial first pass to test our central hypothesis that both copepod density and infection burdens would be linked with deforestation, we used simple Pearson correlation analysis (Fig. 3).

**Figure 3:**
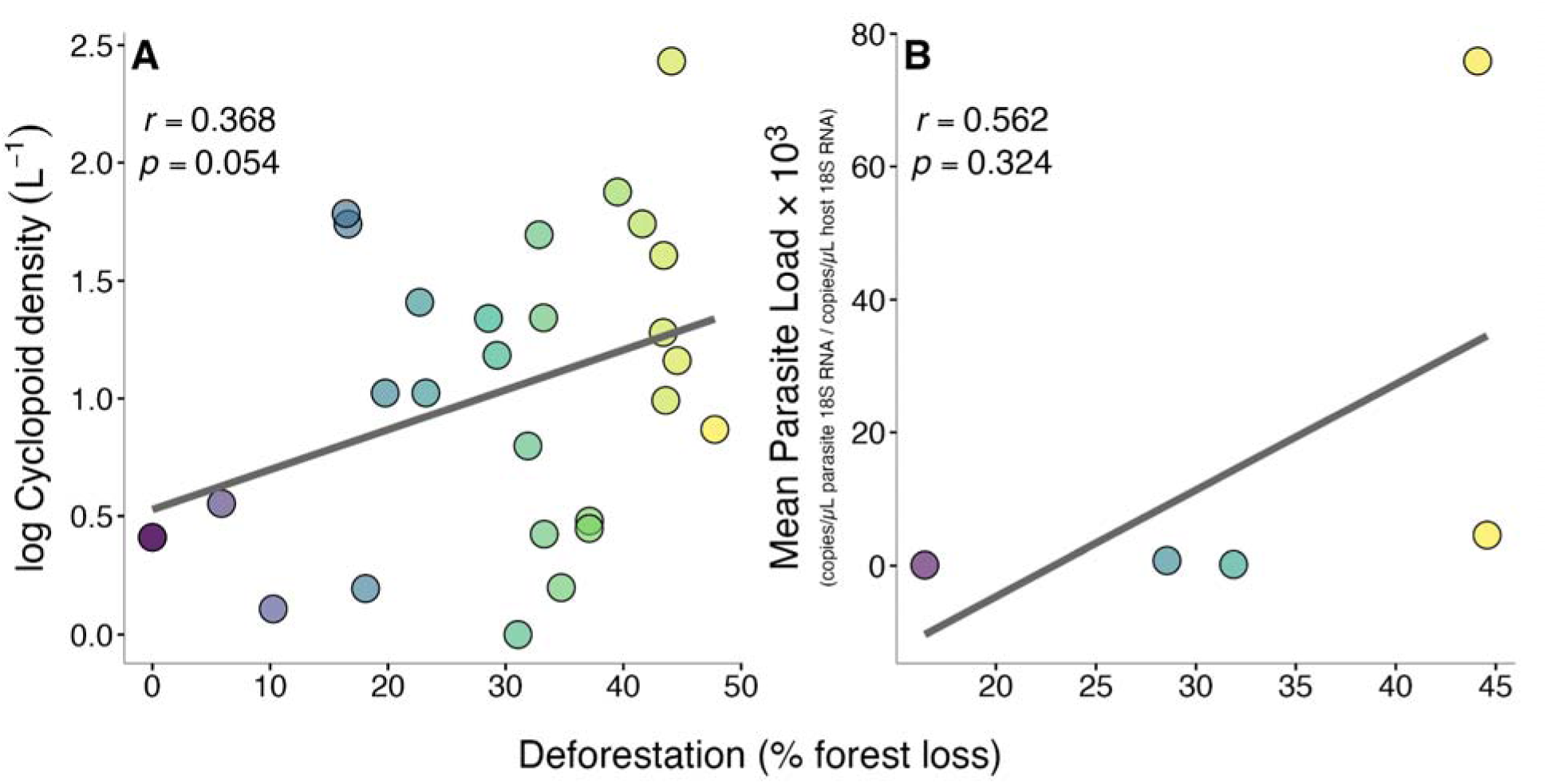
Deforestation is strongly linked to copepod density, but has weak ties with mean parasite load. **(A)** Cyclopoid density (adults and juveniles; excluding nauplii) is positively associated with increasing deforestation (% forest loss), as indicated by the Pearson correlation coefficient *r.* Each point represents the mean copepod density from each of the lakes sampled across Vancouver Island, British Columbia, where we had data on deforestation post-exclusion of non-comparable/outlier lakes (n = 28; of the 39 lakes used for analyses). Deforestation data are proprietary and therefore not available on public databases, making it a challenge to obtain for all surveyed lakes. **(B)** Highly sensitive droplet digital PCR (ddPCR) enabled sensitive detection of extremely low parasite loads within copepod costs. Each point represents a lake on Vancouver Island, British Columbia, that tested positive and had deforestation data available (n = 5; of the 8 lakes that tested positive). Given these low numbers, combined with limited deforestation data, extracting any meaningful patterns with deforestation and infection load is beyond the scope of the current study.

Then, to explore the direct and indirect factors shaping these relationships, we used piece-wise Bayesian structural equation models developed using brms^78,79^ with a Stan backend^80^ via rstan^81^. We quantified hypothesized causal pathways linking percent forest loss (%) to water quality, zooplankton community diversity, copepod host populations, and infection load within hosts using a Bayesian structural equation modeling framework implemented as a manual piecewise SEM (brms R package). Because deforestation data that are released (or not) from logging companies were missing for a substantial fraction of lakes (available for 28 of 39 lakes in this dataset), fitting a single, fully integrated SEM would have forced a complete⍰case analysis and greatly reduced sample size and statistical power across all pathways. To retain an SEM framework while using all available information, we fit the SEM as a piecewise Bayesian SEM, estimating each component equation on the largest subset of lakes with the variables required for that equation. This SEM approach preserves power for pathways not involving deforestation while still allowing land⍰use effects to be evaluated where data exist.

Water quality metrics were integrated over the whole water column depth profiles (integrated chlorophyll a, integrated temperature, integrated dissolved oxygen, and integrated pH). Thermocline depth was calculated from temperature-depth profiles using the thermo.depth function in the rLakeAnalyzer R package^82^. To quantify the extent of deforestation (percent forest loss) was represented by a derived proxy computed from crown closure (deforestation = max(0, 100 - crown_closure)) and is reported as “deforestation (% percent forest loss)”. To stabilize variance for density-like outcomes and support Gaussian likelihoods, chlorophyll and zooplankton density variables were transformed using a natural log plus one transformation, ln(1 + x), after setting negative values to zero prior to transformation (implemented in R using the log1p function).

The SEM structure encoded the following data-driven hypothesis based on the literature (Table 1), with all submodels fit as Gaussian regressions with identity link: temperature was modeled as a function of deforestation; chlorophyll a (chl_log1p) was modeled as a function of deforestation; dissolved oxygen (oxy) was modeled as a function of temperature and chlorophyll a; pH was modeled as a function of chlorophyll; calanoid density (cal_log1p) was modeled as a function of oxygen and pH; and cyclopoid density (cyc_log1p) was modeled as a function of pH and calanoid density.

Each component model was fit with 4 Markov chains with 4,000 iterations per chain (1000 warmup), using rstan as the backend and NUTS control settings adapt_delta = 0.99 and max_treedepth = 15. For each equation, we constructed an equation-specific model data frame and excluded only lakes with missing values in the response or required predictors for that equation, yielding submodel-specific sample sizes.

Direct, indirect, and total effects were computed from posterior draws to propagate uncertainty through the SEM. Direct effects for each arrow were taken as the posterior distribution of the corresponding slope coefficient and summarized by posterior median and 95% credible interval. Indirect effects were computed draw-by-draw as products of coefficients along each simple directed path. Total effects were computed draw-by-draw as the direct effect plus the sum of indirect-path effects for each source-target pair.

To facilitate comparison among paths, standardized effects were computed using global standard deviations for each variable. Coefficients were standardized by rescaling slopes using the predictor and response standard deviations, yielding effects in standard deviation units. The SEM path diagram uses standardized posterior medians as edge labels.

Model performance and adequacy were assessed for each submodel using Markov Chain Monte Carlo (MCMC) diagnostics (R-hat and effective sample sizes) and posterior predictive checks (density overlays). We also recorded standard convergence summaries and NUTS diagnostics (counts of divergences and treedepth saturation). We further generated posterior predictive density overlays for each response and posterior predictive PIT diagnostics computed from posterior predictive draws on the fitted data. PIT ECDFs were compared visually to the Uniform(0,1) reference. Conditional effect plots were generated from the fitted submodel corresponding to each response using the conditional_effects function in brms, with observed lake-level data points overlaid.

## Results

### Univariate Analyses

Cyclopoid copepod density (adults and juveniles; excluding nauplii) and deforestation (percent forest loss) were positively correlated (Fig. 3A; Pearson correlation, r = 0.479, p = 0.018, n = 24). For field data, the strength of this relationship is notable. However, connecting these patterns to changes in within-host infection loads was more challenging. We detected positive infections in eight lakes but only five of these lakes had accompanying deforestation data (Fig. 3B; Pearson correlation, r = 0.562, p = 0.324, n = 5). Due to the high number of true zeros, we limited the statistical analyses here to only the positive lakes and did not use zero-inflated models. Given these low numbers, combined with limited deforestation data, extracting any meaningful patterns with deforestation and infection load is beyond the scope of the current study. Hence, we focus the remainder of our analyses on understanding the direct and indirect pathways linking copepod density and percent forest loss.

### Bayesian Piece-Wise Structural Equation Models

Overall, the SEM indicates that deforestation impacted lake ecosystems through small shifts in water quality captured here by primary production, oxygen, and pH (Fig. 4). However, the full model suggests that quantifying the direct and indirect effects linking deforestation and copepod density remains challenging. Direct effects provided strong support for pathways involving deforestation and water quality. Deforestation was positively associated with chlorophyll a on the ln(1 + x) scale (median 0.00481; 95% credible interval 0.000129 to 0.00947). Oxygen increased with temperature (median 1.59; 95% credible interval 0.275 to 2.93) and decreased with chlorophyll a (median −24.7; 95% credible interval −39.8 to −9.64). Higher chlorophyll a was also associated with lower pH (median −1.24; 95% credible interval −2.03 to −0.444). In contrast, the direct effect of deforestation on temperature was not credibly different from zero (median −0.0508; 95% credible interval −0.115 to 0.0146), and downstream direct relationships from oxygen and pH to calanoids and from pH and calanoids to cyclopoids were imprecisely estimated with 95% credible intervals spanning zero.

**Figure 4:**
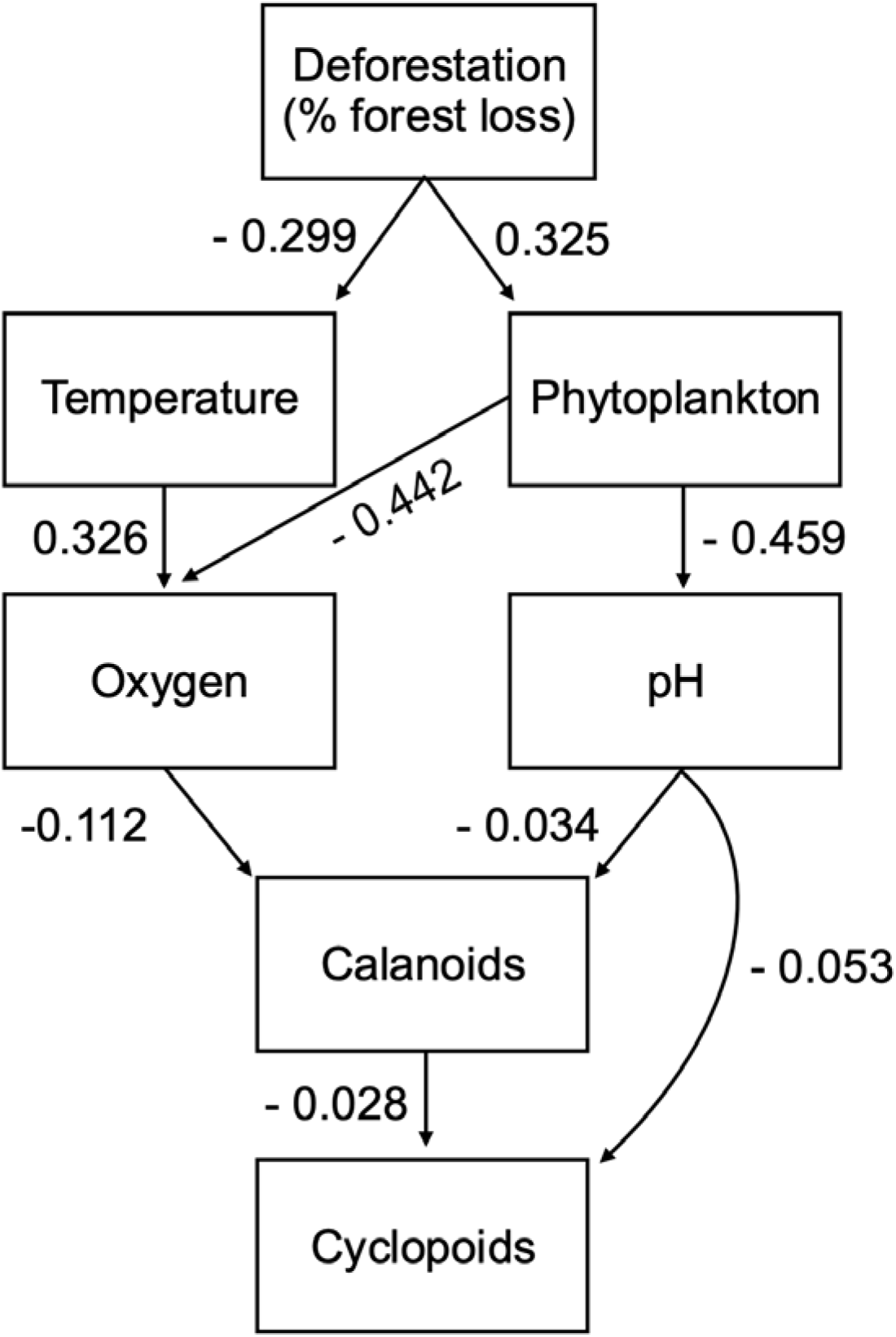
Initial full piecewise Bayesian structural equation model illustrating links between deforestation, water quality, a key competitor, and copepod hosts across 39 lakes on Vancouver Island, British Columbia. The SEM indicates that deforestation is *linked with* predictable shifts in water quality, including higher chlorophylllJ*a* and subsequent reductions in dissolved oxygen and pH. These waterlJquality pathways showed the strongest support in the model, whereas downstream associations involving calanoid and cyclopoid copepods were weaker and estimated with high uncertainty, with 95% credible intervals spanning zero. Indirect effects revealed small negative influences of deforestation on oxygen and pH via increases in chlorophyll *a*, but total effects of deforestation and chlorophyll *a* on copepod densities were not credibly different from zero. Note: Unlike traditional covariancelJbased SEMs that report standardized correlation coefficients, the model framework used here presents standardized posterior median *total effects* for each path (direct + indirect), providing an integrated measure of association across modeled pathways. Arrow labels are standardized posterior medians based on total effects for each source-target pair.

Indirect effects highlighted the cascading consequences of deforestation mediated through changes in chlorophyll a (a proxy for phytoplankton biomass, which typically increases with nutrient inputs). Deforestation had a negative indirect effect on dissolved oxygen via chlorophyll a (deforestation → chlorophyll a → oxygen; median −0.113; 95% credible interval −0.277 to −0.00231) and a negative indirect effect on pH via chlorophyll a (deforestation → chlorophyll a → pH; median −0.00556; 95% credible interval −0.0142 to −0.0000693). Consistent with these indirect pathways, the total effect of deforestation on dissolved oxygen was negative (median −0.233; 95% credible interval −0.491 to −0.0379), whereas the total effects of chlorophyll a on calanoids and cyclopoids, and of deforestation on calanoids and cyclopoids, were not credibly different from zero.

While this particular SEM framework is designed to address the limitations inherent in complex field data, missing data components, and low sample sizes, the ability to quantify the effects of multiple stressors remains challenging. Therefore, we also developed an additional model that compiled each water quality factor into a single variable. Specifically, we included chlorophyll a, dissolved oxygen, pH, and temperature as a z-scored composite variable for water quality. With this reduced model, we were also able to include a direct path from deforestation to cyclopoids.

Overall, the reduced model indicates that deforestation had a positive total effect on cyclopoid density (median 0.361; 95% credible interval -0.0407 to 0.755). None of the individual pathways from deforestation to cyclopoids showed strong directional evidence on their own, whether through direct effects or through indirect routes involving water⍰quality changes or shifts in other zooplankton groups. However, when all pathways were considered together, the model indicated that deforestation was linked with increased cyclopoid densities through a series of small, incremental changes that accumulated across the system. In other words, while each individual connection was modest, their combined influence produced a clear overall effect (Fig. 5).

**Figure 5:**
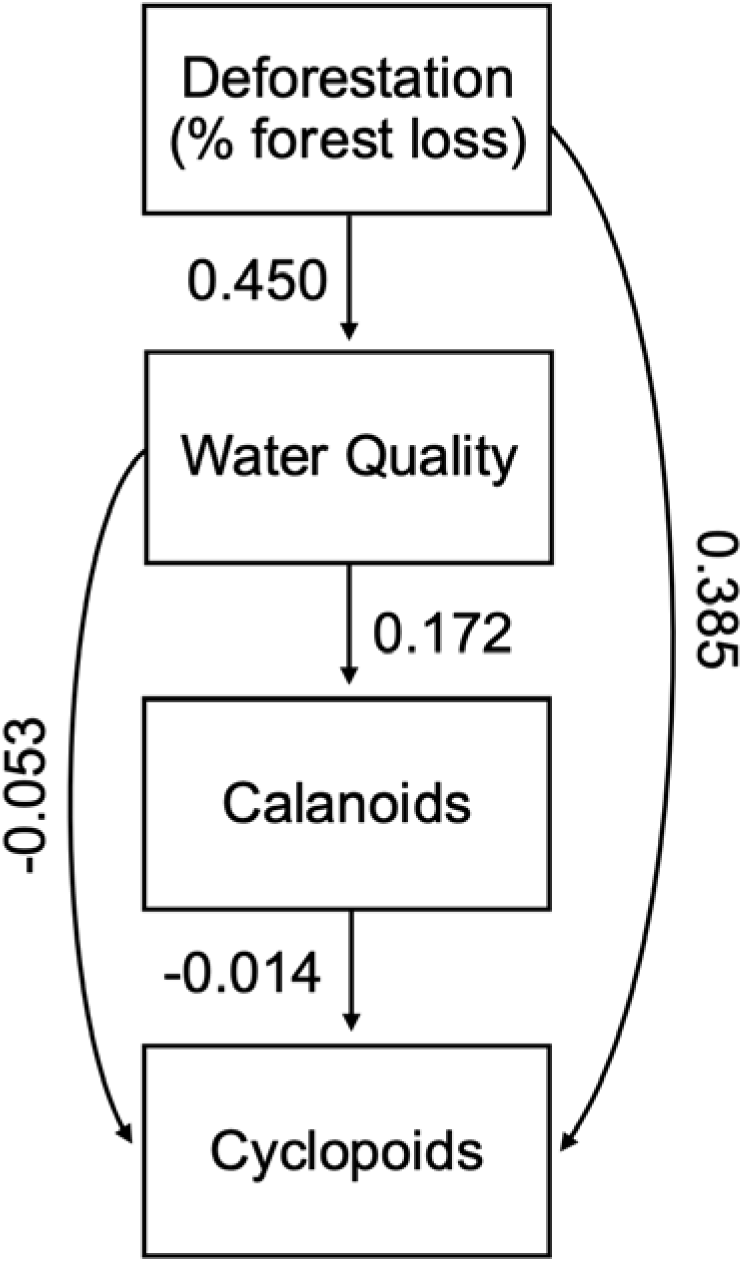
Piecewise Bayesian structural equation model illustrating links between deforestation, water quality, a key competitor, and copepod hosts across 39 lakes on Vancouver Island, British Columbia. Each submodel was fitted on a subset of lakes with a complete dataset for response and predictor variables: calanoid (n = 39), water quality (n = 28), and cyclopoid (n = 28). The water quality variable is a z-scored composite of chlorophyll, temperature, oxygen, and pH. This reduced SEM model shows that deforestation had a positive total effect on copepod density. No single pathway from deforestation to cyclopoids exhibited strong directional influence on its own, whether through direct effects or via indirect routes involving waterlJquality changes or shifts in other calanoids. However, when all pathways were considered together, the model revealed that deforestation was linked with higher cyclopoid densities through a series of small, cumulative changes of small effect. Thus, while each individual link was modest, their combined influence produced a more pronounced overall effect. Note: Unlike traditional covariancelJbased SEMs that report standardized correlation coefficients, the model framework used here presents standardized posterior median *total effects* for each path (direct + indirect), providing an integrated measure of association across modeled pathways. Arrow labels are standardized posterior medians based on total effects for each source-target pair.

Deforestation did not have a direct effect on cyclopoid density that was credibly different from zero (median 0.0181; 95% credible interval -0.00558 to 0.0412) but had a stronger positive association with changes in water quality (median 0.0328; 95% credible interval 0.0158 to 0.0497). Water quality was weakly associated with cyclopoid (median -0.0339; 95% credible interval -0.478 to 0.413) and calanoid density (median 0.0929; 95% credible interval -0.0857 to 0.270). The direct effect of calanoids on cyclopoids was not credibly different from zero (median -0.0169; 95% credible interval -0.667 to 0.651).

None of the indirect effects were credibly different than zero. Deforestation was linked to cyclopoid density through two pathways: shifts in water quality (deforestation → water quality → cyclopoid density; median -0.00102; 95% credible interval -0.00166 to 0.0143), and calanoid density (deforestation → water quality → calanoid density → cyclopoid density; median - 0.0000147; 95% credible interval -0.00316 to 0.00295). Water quality was associated with cyclopoid density through shifts in calanoid density (water quality → calanoid density → cyclopoid density; median -0.000489; 95% credible interval -0.936 to 0.0891). Additionally, deforestation was linked to calanoid density via changes in water quality (deforestation → water quality → calanoid density; median 0.00284; 95% credible interval -0.00282 to 0.00976).

## Discussion

Our results reveal an intriguing relationship between deforestation and copepod density, with densities increasing with decreasing forest cover. We further show that this pattern is likely mediated by deforestation-driven changes in water quality. Lakes impacted by greater percent forest loss exhibited higher phytoplankton biomass and elevated pH, both consistent with nutrient enrichment. Such conditions are known to favor cyclopoids across freshwater systems, where they are positively associated with enrichment indicators, including chlorophyll a^29,40,41,44^. However, our results suggest that while deforestation reliably reshapes lake water quality, its influence on copepod densities likely operates through additional ecological pathways not captured here, underscoring the complexity of linking land⍰use change to disease dynamics in natural ecosystems.

Altogether, the results suggest that deforestation’s influence on copepod density arises from a broader suite of ecological cascades, some captured in our SEM, others operating outside the pathways we measured. This mismatch between strong system⍰level correlations and weak quantitative capacity for indirect effects highlights both the strengths of SEMs for identifying where ecological linkages break down and the inherent complexity of attributing biotic responses to individual drivers in multi⍰stressor landscapes^83–86^. These broader, unresolved pathways align with additional, likely non-linear, mechanisms through which land⍰use change can restructure zooplankton communities.

For instance, multiple interlinked abiotic and biotic drivers (Table 1) could mediate land-use driven changes to cyclopoid populations, creating complex, non-linear, feedback loops. Apart from direct increases in phytoplankton, nutrient-enriched lakes also tend to have lower dissolved oxygen levels due to rapid plankton decomposition^87–89^. This can favor cyclopoids, partly due to greater degrees of hypoxia tolerance as compared to larger-bodied zooplankton and zooplanktivores, potentially alleviating both competitive and predation pressures stifling cyclopoid population growth^27,90–92^. Lake depth is an additional factor to consider, with deeper lakes supporting greater populations of larger-bodied zooplankton, sustaining competing zooplankton species that are otherwise subject to significant predation in shallow lakes, where cyclopoids dominate due to their relatively smaller size^41,42,93–95^.

Additionally, land-use effects may also operate indirectly by reshaping zooplankton community composition, with shifts towards copepod-dominated communities, and away from cladoceran-dominated ones^11,43,92,93,96–99^. Such shifts in community composition alter grazing pressure, energy transfer, and food web structure, in ways that can modify infectious disease dynamics^1,4,5,36,100–106^. These consequential changes in infection dynamics are likely driven by nonlinear feedback that is difficult to capture with limited sample sizes and current statistical methods. Future studies that investigate such nonlinear, cross-scale, and bidirectional feedback through simulation models can help guide future empirical work.

Links between land use change and infectious disease have only recently emerged as key ecological levers to prevent zoonotic spillover and mitigate the (re)emergence of outbreaks. Such connections may be particularly relevant for the numerous zoonotic diseases associated with copepods, which serve as the first intermediate host for multiple macro- and microparasites, with consequences for both agricultural systems and human health^22,31^. Their sensitivity to land⍰use change makes them especially responsive to the kinds of environmental shifts described above, raising the possibility that changes in copepod populations may serve as early indicators of broader ecological risk. Given their pivotal role in aquatic food webs, we asked whether copepods could also serve as indicators of disease risk for macroparasites in increasingly human⍰modified ecosystems.

Addressing this question has historically been constrained by the difficulty of detecting parasites within copepod hosts. Conventional approaches rely on tissue preservation followed by labor-intensive microscopy, which often fails to identify early stages of infection and can degrade parasite tissues or DNA^107,108^. These limitations have hindered efforts to evaluate copepods as indicators of parasite risk despite their central role in transmission pathways. Common molecular approaches, such as qPCR, often struggle to reliably quantify low-abundance DNA^107–109^. This limitation is particularly acute in understudied helminth systems, where detecting infections in invertebrate intermediate hosts requires analytical methods with high sensitivity.

To overcome these constraints, we employed droplet digital PCR (ddPCR) to design a multiplexed assay to reliably detect infection in cyclopoids at concentrations as small as 0.1 picogram of sample DNA^110^. ddPCR allows for high-throughput quantification of target gene concentrations, independent of the standard curves required with other quantification methods^111^. This is particularly powerful when investigating understudied parasites, where generating a standard curve may be logistically infeasible. Additionally, ddPCR analytics also enable accurate detection of DNA at much lower concentrations, and capture target DNA spanning a wider gradient^110,111^.

Yet, even with this highly sensitive approach, infected copepods were detected in only eight of the lakes that we surveyed. This result is unlikely to reflect purely methodological limitations, given that these samples fell within recommended ranges for optimal quantification^110^. Instead, several ecological mechanisms may explain the low detection of infected copepods. Parasite-induced behavioral shifts likely increase predation risk: infected copepods exhibit predator avoidance early in infection but transition to low-motility behaviors once parasites reach the infective stage, making them more susceptible to fish consumption^58,63,112,113^. Selective predation could therefore remove infected individuals from the sampling pool, biasing captures toward uninfected copepods.

Additionally, sampling timing may have further contributed to low detection, as collections were aligned with seasonal dynamics of the fish host and may have occurred near the end of the optimal window for detecting infected copepods^47,114^. Preliminary data from the subsequent field season support this possibility, suggesting that earlier summer sampling captures more infected individuals (Srinivas et al., unpublished data). More broadly, the absence of long-term temporal data limits our ability to identify peak infection periods, underscoring the need for longitudinal studies to resolve seasonal transmission dynamics in copepod hosts^115^.

Future steps will focus on examining long-term infection data to identify peak transmission windows and potential ecological mechanisms underlying low detection rates of infected copepods. Quantifying copepod infection variability across depth gradients within a lake, as opposed to using a singular lake-level metric, may resolve weak or inconsistent linkages between water quality metrics, host density, and infection levels^116^. Methodologically, increased ddPCR replication and improved DNA extraction efficiencies could further bolster detection probability. Laboratory experiments to assess how infection affects individual copepods could inform how host-level fitness consequences translate to population-level dynamics^71^. Finally, linking infection metrics across successive hosts with environmental factors and land-use gradients would help elucidate how land-use change affects not only copepod density but also parasite transmission across hosts, providing a more mechanistic understanding of how ecosystem alteration shapes disease risk.

Given the unpredictability of disease outcomes in human-altered landscapes, identifying reliable indicators of parasite transmission is critical, particularly for aquatic ecosystems that are sensitive to anthropogenic stressors like land-use change^1,9,12,15,117^. Copepods occupy a pivotal role in aquatic food webs, and respond strongly to environmental changes, serving as an integral ecological link through which land-use can influence parasite transmission and zooplankton community composition^31,41,43,44,96–98,118^. These factors make copepods an ideal candidate for monitoring land-use driven cascades in host-parasite systems, and to serve as potential early-warning indicators of disease outbreaks. Despite not detecting a strong correlation between land-use and within-host parasite load, our results link copepod host populations to land-use mediated changes in water quality, highlighting a potential mechanistic pathway to understand land-use driven disease dynamics. More broadly, by connecting copepod population responses to land-use change, not only does this study inform host-parasite biology, but it also provides a potential ecological lever to manage infectious diseases in increasingly human-altered landscapes.

## Land Acknowledgement

We respectfully acknowledge that this research was conducted across sites situated within traditional territories of Nuu-chah-nulth, Kwakwaka’wakw, and Laich-kwil-tach nations on Vancouver Island, British Columbia. We acknowledge their long and proud history in their traditional territories.

## Author Contributions

Conceptualization: DIB, AKH, JLH. Developing Methods: DIB, AKH, JLH, POS, JB, CAF, HA, ENE, GV, EC, IS. Data Analysis: JLH, POS, IS. Preparation of figures and tables: POS, AKH, JLH, JB, IS. Writing: JLH and IS wrote the first draft; all authors contributed to revisions. Conducting the research, data interpretation: DIB, AKH, JLH, POS, JB, CAF, ENE, HA, EC, GV, IS, SD, CP, GS, AC.

## Data Availability

All data and code supporting the findings of this study will be deposited in the Environmental Data Initiative repository upon acceptance for publication. Data and associated analysis code are currently hosted at GitHub (https://github.com/jhite-eco-epi/Landuse-water-quality-copepods) and will be archived in a permanent repository prior to publication. Code used for geospatial modeling is available at GitHub (https://github.com/jberini/vancouver-island-amr). Due to size and complexity, geospatial datasets are available upon request.

