## Supplementary Information for "Linking land□use change, water quality, and host-parasite dynamics with droplet digital PCR and Bayesian path analyses"

Hund: 0000-0001-7758-6757

Bolnick: 0000-0003-3148-6296

Hite: 0000-0003-4955-0794

**Keywords:** Deforestation, helminths, macroparasites, trophic transmission, copepods

**Supporting Information Text 1 (Methods)**

*Zooplankton Community Composition:*

To identify the zooplankton community present in each lake, within a lake sample, copepods were categorized into *Calanoida spp*., *Cyclopoida* *spp*., and nauplii (irrespective of genus). For cladocerans, the following taxa were identified: *Daphnia spp*., *Bosmina spp*., *Holopedium spp*., *Chydorus spp*., *Diaphanosoma spp*., and the family Polyphemidae. Each lake sample was diluted to 100 mL. A maximum of five subsamples (2 mL each) and a minimum of two subsamples were taken from each 100 mL dilution without replacement. For a given lake, a maximum of n = 100 copepods, and n = 30 for non-copepod zooplankton, were counted. If, after counting and identifying all the members within the first two subsamples, we determined that further subsampling was unlikely to result in meeting count aims, no further subsamples were taken.

To obtain densities of zooplankton across lakes, the count data were divided by the sampled water volume. The volume of water filtered per tow was calculated as the surface area of the tow x tow depth. Densities were log-transformed (log(1+x)) prior to analysis. Subsequently, for each lake, the ratio of calanoids to cyclopoids, as well as the overall ratio of copepods to cladocerans, was calculated.

**Supporting Information Text 2 (Methods)**

*Univariate Analysis*

We also examined relationships between deforestation, water quality, copepod density, and parasite infection across the 39 lakes (of the 47 sampled; post-exclusion), using Pearson's correlation analysis. All statistical analyses were conducted in R version 4.4.2^72^ and were performed at the lake level. Figures were generated in R using ggplot2^73^, with multi-panel layouts assembled using patchwork^74^.

**Supporting Information Text 3 (Results)**

*Results*

To identify potential proximate abiotic drivers mediating the relationship between copepod density and percent forest loss, we explored our hypothesized relationships (Table 1) between percent forest loss and water quality metrics (Fig. S1), and subsequent changes in copepod density associated with variation in water quality (Fig. S2). We also examined relationships between land-use change and zooplankton community composition to assess whether copepods increased disproportionately relative to other zooplankton taxa with increasing percent forest loss (Fig. S3). Lastly, we explored how water quality metrics were associated with parasite infection across the subset of infected lakes (Fig. S4).

**Water quality change potentially mediates the association between deforestation and copepod density.** Our exploration of the relationships between percent forest loss and water quality metrics reveals that, as predicted, phytoplankton (log_10_ Chlorophyll-*a* RFU) - which typically increases on nutrient enrichment - was positively associated with percent forest loss (Fig. S1A; Pearson correlation, *r* = 0.388, *p* = 0.041). Water temperature decreased with percent forest loss, which did not support our prediction that reduced riparian shading should increase water temperature (Fig. S1B; Pearson correlation, *r* = -0.297, *p* = 0.125). Decreasing lake pH was significantly linked to increasing percent forest loss (Fig. S1C; Pearson correlation, *r* = -0.729, *p* < 0.001), which may potentially be mediated by increases in phytoplankton. Not surprisingly, the relationship between percent forest loss and dissolved oxygen was negative (Fig. S1D; Pearson correlation, *r* = -0.292, *p* = 0.132).

Copepod density was weakly linked to increases in phytoplankton (log_10_ Chlorophyll-*a* RFU; Fig. S2A; Pearson correlation, *r* = 0.059, *p* = 0.72) and dissolved oxygen (Fig. S2D; Pearson correlation, *r* = 0.063, *p* = 0.703), although this may be due to most sampled lakes having a mean dissolved oxygen level > 10 mg L^-1^. Interestingly, in contrast to our predictions, increasing water temperature was associated with a decrease in copepod density (Fig. S2B; Pearson correlation, *r* = -0.205, *p* = 0.211). Additionally, increasing pH was weakly linked to decreasing copepod density (Fig. S2C; Pearson correlation, *r* = -0.057, *p* = 0.73).

**Deforestation is linked to shifts in zooplankton community composition, potentially leading to copepod-dominated communities.** The ratio of Cladocera:Copepoda decreased with greater percent forest loss (Fig. S3A; Pearson correlation, *r* = -0.281, *p* = 0.14), indicating a shift from Cladocera-dominated to copepod-dominated communities with increasing percent forest loss. Similarly, the ratio of Calanoida:Cyclopoida was weakly linked to increasing percent forest loss, indicating a shift from larger-bodied Calanoida to smaller-bodied cyclopoid copepods (Fig S3B; Pearson correlation, *r* = -0.411, *p* = 0.033). Additionally, a correlation matrix also shows that broadly, changes in cyclopoid density could indicate cascading shifts in zooplankton community composition, with deforestation potentially causing a decline in the diversity of functional zooplankton groups (Fig. S3C).

**Water quality metrics are weakly associated with mean parasite load.** Mean parasite load was negatively linked with increasing phytoplankton (log_10_ Chlorophyll-*a* RFU; Fig. S4A; Pearson correlation, *r* = -0.027, *p* = 0.095), and lake temperature (Fig. S4B; Pearson correlation, *r* = -0.223, *p* = 0.63). pH had a stronger negative link to mean parasite load (Fig. S4C; Pearson correlation, *r* = -0.618, *p* = 0.139). Additionally, increased dissolved oxygen was weakly associated with a decline in mean parasite load (Fig. S4D; Pearson correlation, *r* = -0.036, *p* = 0.939). Interestingly, we explore two additional metrics of nutrient enrichment - dissolved organic matter (log_10_ fDOM RFU) and cyanobacteria (log_10_ Phycocyanin RFU) - with the latter showing a stronger association with mean parasite load than phytoplankton (Fig. S4E-F). Given the small subset of lakes with infected copepods, this suggests that in the presence of stronger infection signals, distinct resource-mediated links between land-use change and infection patterns could emerge.

**Supporting Information Figures (Results)**

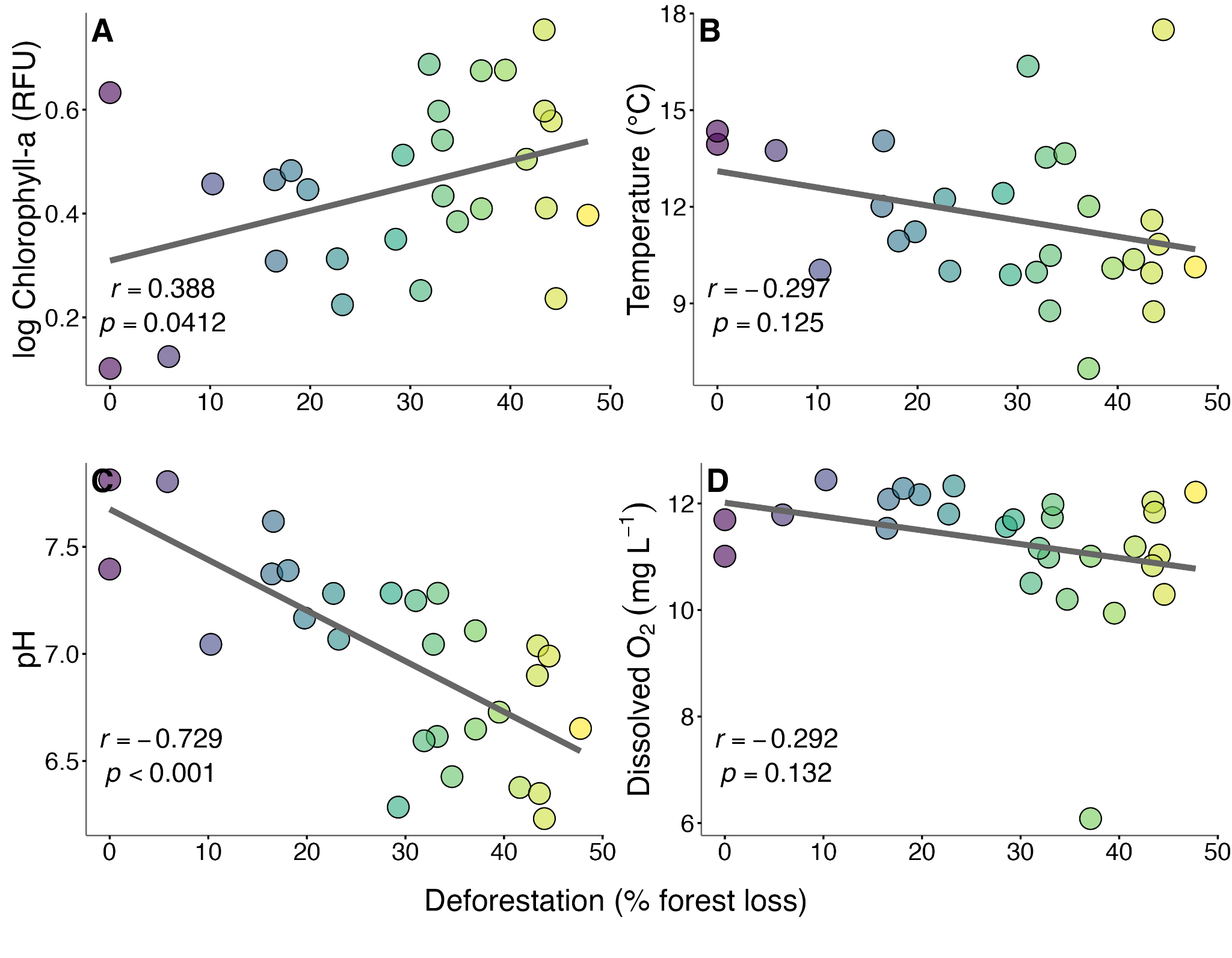

**Figure S1:** **Links between deforestation and water quality could potentially help explain shifts in cyclopoid abundance.** Each point shows the mean vertical profile for an individual lake (n = 28). **(A)** Phytoplankton (log_10_ Chlorophyll-*a* RFU) shows a positive association with increasing percent forest loss (%), while **(B)** water temperature (℃), **(C)** pH, and **(D)** Dissolved oxygen (DO; mgL^-1^) show a negative association with increasing forest loss.

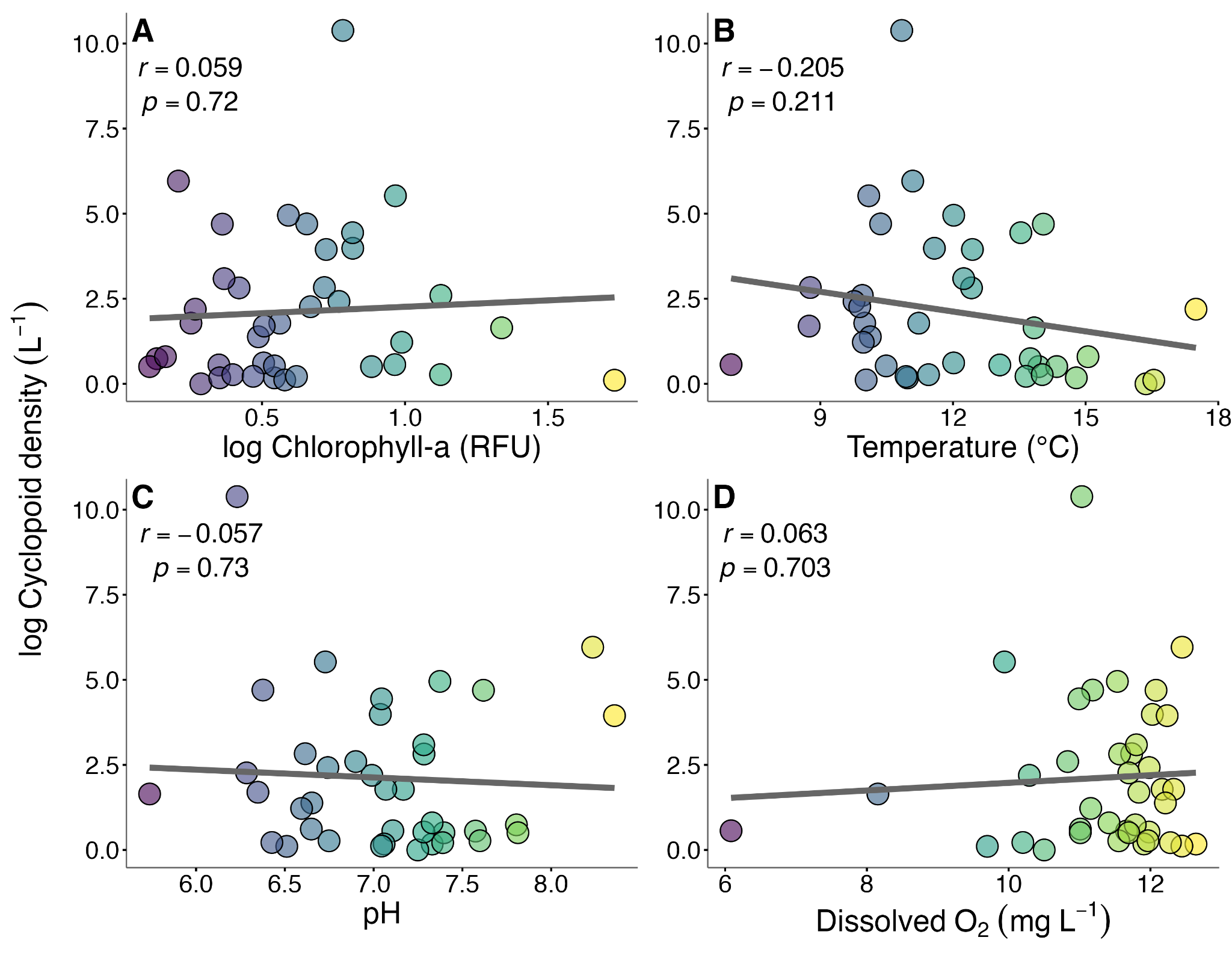

**Figure S2:** **Cyclopoid density is linked to water quality metrics, highlighting potential mechanisms by which percent forest loss could shape copepod populations.** Each point shows the mean vertical profile for an individual lake (n = 39). **(A)** Phytoplankton (log_10_ Chlorophyll-*a* RFU) shows a weak positive relationship with cyclopoid density. **(B)** Water temperature (℃) shows a negative association with cyclopoid density. **(C)** pH shows a weak negative relationship with cyclopoid density. **(D)** Dissolved oxygen (DO; mg L^-1^) shows a weak positive relationship with cyclopoid density.

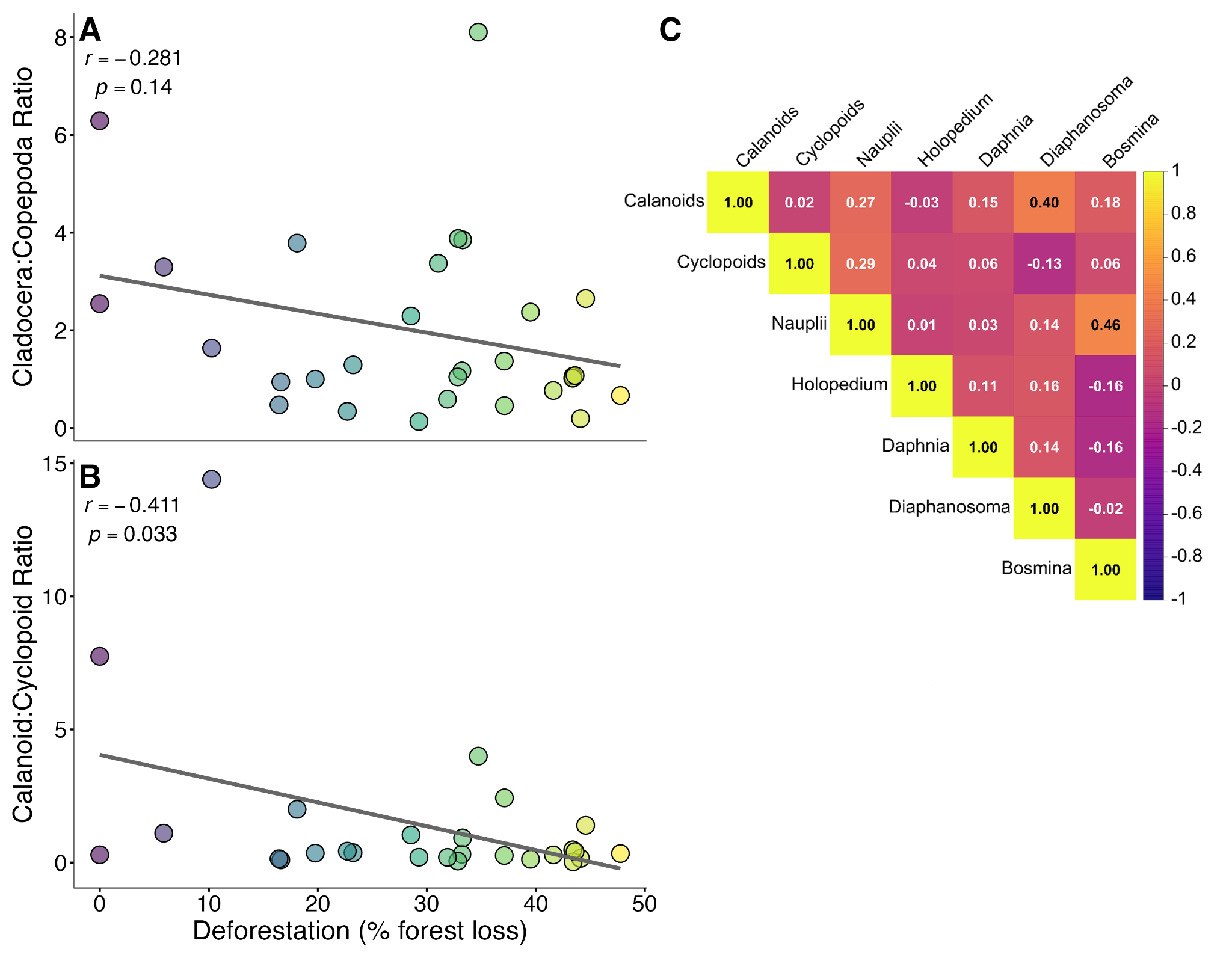

**Figure S3: Variation in percent forest loss is linked to shifts in zooplankton community composition.** Each point shows the mean vertical profile for an individual lake. **(A)** The relative proportion of Cladocera (cladocera:copepoda) in a lake (n = 28) is negatively associated with increasing percent forest loss (%). **(B)** The relative proportion of Calanoida (calanoida:cyclopoid) decreases with increasing percent forest loss (%) across lakes (n = 28). **(C)** Heat map depicting correlations between zooplankton groups observed across lakes (n = 39).

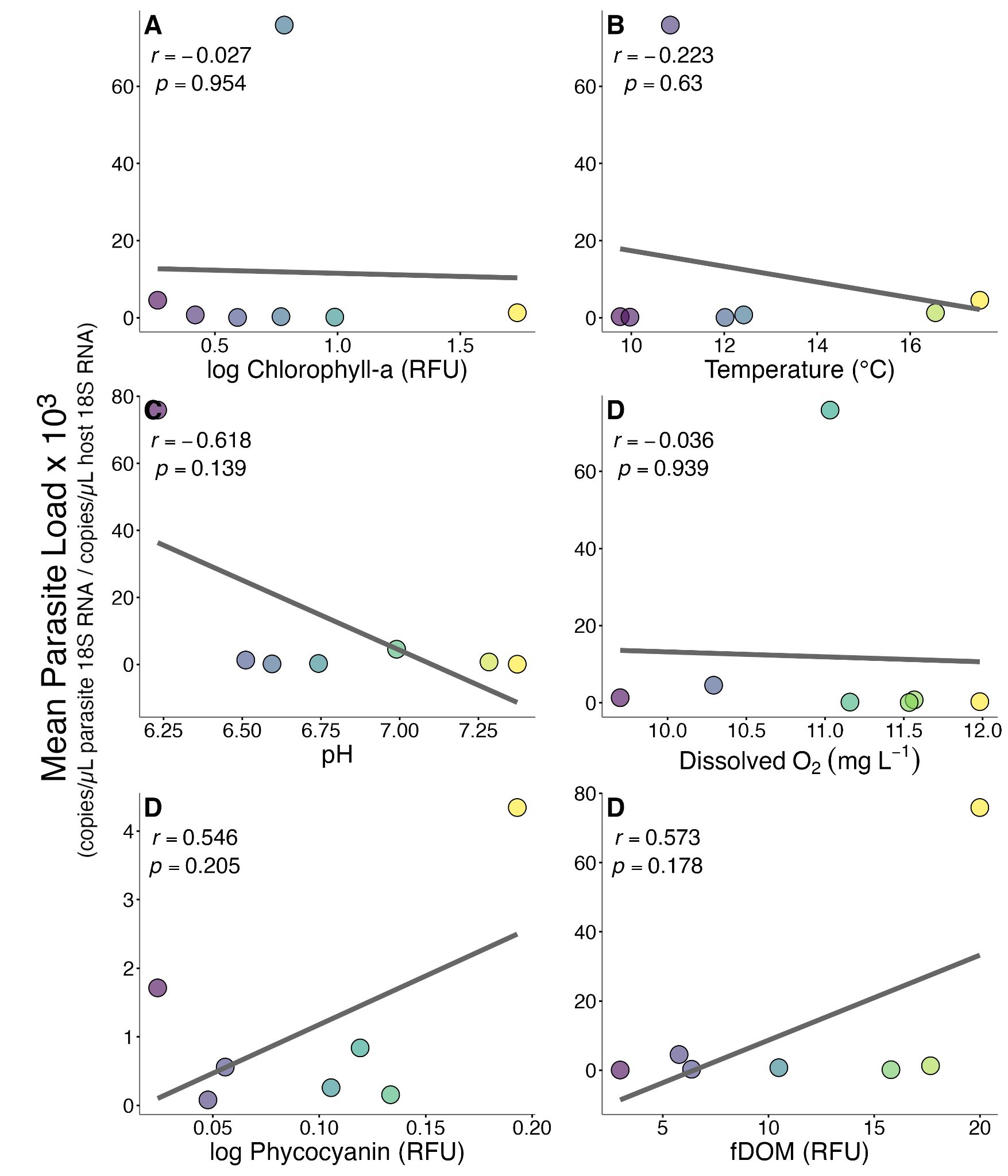

**Figure S4: Mean parasite load is weakly linked to water quality metrics.** Each point shows the mean vertical profile for an individual lake. **(A)** Phytoplankton (log_10_ Chlorophyll-*a* RFU) and **(B)** water temperature (℃) show weak negative associations with infection, while **(C)** lake pH shows a stronger negative association with mean parasite load. **(D)** Additionally, dissolved oxygen (mgL^-1^) is weakly associated with mean parasite load. **(E)** Allochthonous inputs of dissolved organic matter (log_10_ fDOM RFU) and **(F)** Cyanobacteria (log_10_ Phycocyanin RFU) are largely positively associated with mean parasite load.
